# Sensory Stimulation-Induced Astrocytic Calcium Signaling in Electrically Silent Ischemic Penumbra

**DOI:** 10.1101/652305

**Authors:** Reena P. Murmu, Jonas C. Fordsmann, Changsi Cai, Alexey Brazhe, Kirsten J. Thomsen, Martin Lauritzen

## Abstract

Middle cerebral artery occlusion (MCAO) induces ischemia characterized by a densely ischemic focus, and a less densely ischemic penumbral zone in which neurons and astrocytes display age-dependent dynamic variations in spontaneous Ca^2+^ activities. However, it is unknown whether penumbral nerve cells respond to sensory stimulation early after stroke onset, which is critical for understanding stimulation-induced stroke therapy. In this study, we investigated the ischemic penumbra’s capacity to respond to somatosensory input. We examined adult (3- to 4-month-old) and old (18- to 24-month-old) male mice at 2–4 hours after MCAO, using two-photon microscopy to record somatosensory stimulation-induced neuronal and astrocytic Ca^2+^ signals in the ischemic penumbra. In both adult and old mice, MCAO abolished spontaneous and stimulation-induced electrical activity in the penumbra, and strongly reduced stimulation-induced Ca^2+^ responses in neuronal somas (35–82%) and neuropil (92–100%) in the penumbra. In comparison, after stroke, stimulation-induced astrocytic Ca^2+^ responses in the penumbra were only moderately reduced (by 54–62%) in adult mice, and were even better preserved (reduced by 31–38%) in old mice.

Our results suggest that somatosensory stimulation evokes astrocytic Ca^2+^ activity in the ischemic penumbra. We hypothesize that the relatively preserved excitability of astrocytes, most prominent in aged mice, may modulate protection from ischemic infarcts during early somatosensory activation of an ischemic cortical area. Future neuroprotective efforts in stroke may target spontaneous or stimulation-induced activity of astrocytes in the ischemic penumbra.

## Introduction

Stroke patients experience extensive sensory and tactile stimulation during transport and critical care. Experimental studies show potential beneficial effects of early sensory stimulation of peri-infarct tissue (Lay et al., 2011; 2012; Frostig et al., 2013; Hancock et al., 2013; Lay et al., 2013). However, the results are controversial and the mechanisms underlying stimulation-induced neuroprotection remain elusive (Baron, 2018). Immediately following ischemia, glutamate accumulates at synapses, (Drejer et al., 1985) resulting in extensive stimulation of N-methyl-D-aspartate (NMDA) receptors that can eventually become neurotoxic (Tu et al., 2010; Lai et al., 2014; Chamorro et al., 2016). Unfortunately, little is presently known about astroglial function in the penumbra and in neuroprotection. These cells perform important functions, including neurotransmitter uptake and recycling, (Schousboe, 2018) neurovascular coupling, (Lind et al., 2018) and blood-brain barrier maintenance (Kutuzov et al., 2018). In the current study, we aimed to elucidate the evoked activity of neurons and astroglial cells in the penumbra, to better understand the mechanisms underlying somatosensory stimulation-induced stroke protection.

Ischemic strokes are more common among the elderly than in the young population; therefore, translation of preclinical results to clinical stroke requires inclusion of an aged animal group (O’Collins et al., 2006; Dirnagl and Macleod, 2009; Chen et al., 2010; Chisholm and Sohrabji, 2016). Our present study included both adult and aged mice. While electrical signals are silenced in the ischemic penumbra, both neurons and astrocytes produce spontaneous Ca^2+^ activity. We previously demonstrated that this relies on the preservation of neuronal excitability, which reflects some degree of functional organization (Fordsmann et al., 2018). Importantly, adult mice show reduced spontaneous Ca^2+^ activities (astrocytic and neuronal), whereas old mice exhibit unchanged spontaneous neuronal Ca^2+^ activity and abundant astrocytic Ca^2+^ activity that is modulated by synaptic activity (Fordsmann et al., 2018). Notably, intracellular Ca^2+^ changes represent biochemical signals and are not reflected in electrical signals.

Here we examined the possibility that somatosensory stimulation evoked neuronal and astrocytic Ca^2+^ responses in the ischemic penumbra despite electrical silence. We examined adult and old male mice at 2–4 hours after middle cerebral artery occlusion (MCAO), using a combination of two-photon Ca^2+^ imaging, laser speckle imaging of cerebral blood flow, and electrophysiological recordings. In both adult and old mice, stroke strongly reduced stimulation-induced Ca^2+^ responses in neurons, while stimulation-induced astrocytic Ca^2+^ responses were preserved. This suggests that somatosensory stimulation of the electrically silent ischemic penumbra induced increased astrocyte Ca^2+^ within a time-window that is relevant for early stroke therapy.

## Materials and Methods

### Animal Handling

This study included 42 male C57Bl/6 mice (Janvier labs): 21 adult and 21 old. In each mouse, the trachea was cannulated for mechanical ventilation (SAAR-830; CWE), and a catheter was inserted into the left femoral artery to monitor blood pressure and blood gases. Another catheter was placed in the femoral vein for anesthetic infusion (α-chloralose). To ensure that the mice were maintained under physiological conditions, we continuously monitored end-expiratory CO_2_ (Capnograph type 340; Harvard Apparatus) and blood pressure (Pressure Monitor BP-1; World Precision Instruments). Arterial blood gases were repeatedly measured (PO_2_, 95–110 mmHg; PCO_2_, 35–40 mmHg; pH, 7.35–7.45; ABL 700Series Radiometer). Body temperature was maintained at 37°C using a rectal temperature probe and a heating blanket (Model TC-1000 Temperature Controller; CWE).

All procedures involving animals were approved by the Danish National Ethics Committee (#2014-15-0201-00027), and were performed in accordance with the guidelines outlined in the European Council’s Convention for the Protection of Vertebrate Animals used for Experimental and Other Scientific Purposes, and in compliance with the Animal Research: Reporting of *In Vivo* Experiments (ARRIVE) guidelines.

Part of the dataset for the 42 mice examined by two-photon microscopy imaging has previously been described (Fordsmann et al., 2018). In our previously published study, we report spontaneous Ca^2+^ changes using a different analytical method than in the present study. We did not previously describe stimulation-induced changes in Ca^2+^ signaling, and the reported two-photon imaging data do not overlap between the two manuscripts.

### Anesthesia

Anesthesia was induced with intraperitoneal bolus injections of the α2-adrenergic-receptor agonist xylazine (10 mg kg^−1^) and the NMDA-receptor antagonist ketamine (60 mg kg^−1^). To maintain anesthesia during surgery, we administered supplemental doses of ketamine (30 mg kg^−1^/20 min). Anesthesia level was assessed by testing the motor response to a hind-limb pinch. For the duration of the experiment, the mice were supplied with NaHCO_3_ (200 mM, 0.15 ml/2 h, i.p.) to prevent metabolic acidosis and declining blood pressure. Upon completion of all surgical procedures, anesthesia was switched to continuous intravenous infusion with α-chloralose (25 mg/kg/h, i.v.) (Figure 1).

**FIGURE 1:**
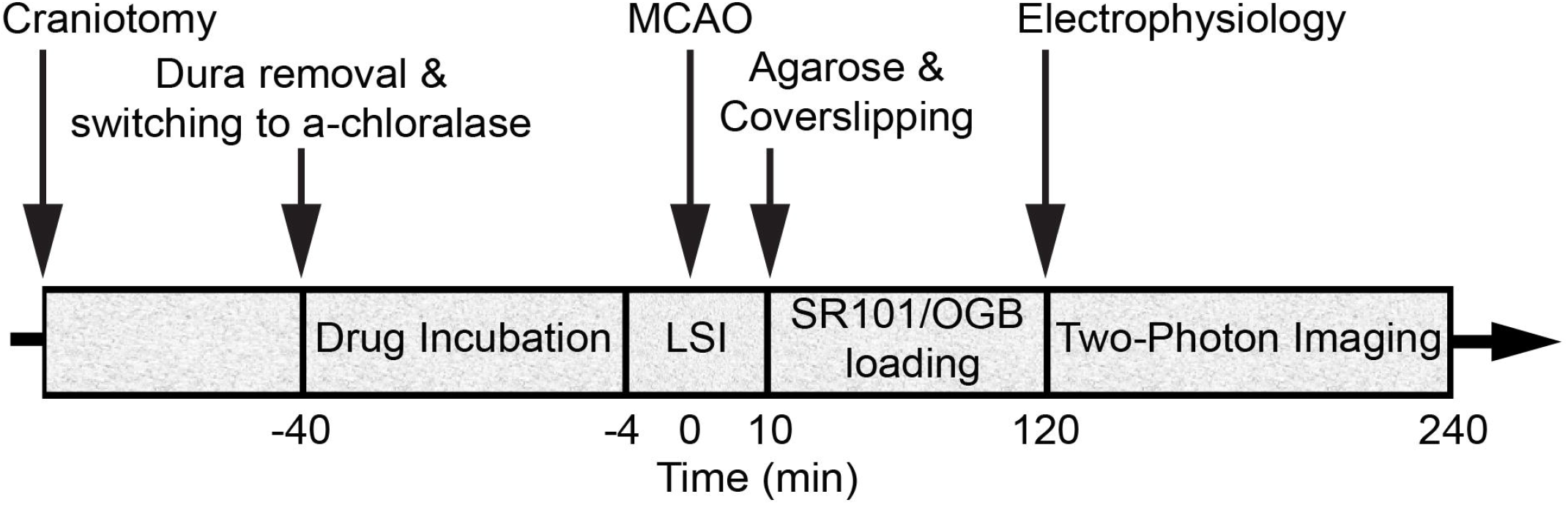
Diagram of the experimental timeline. Middle cerebral artery occlusion (MCAO) was induced during laser speckle imaging (LSI) of cerebral blood flow (CBF). Extracellular local field potentials (LFPs) were recorded using a single-barreled glass micropipette that contained the Ca^2+^-sensitive dye Oregon Green 488 BAPTA-1/AM (OGB) and the astrocyte-specific marker sulforhodamine 101 (SR101).

During surgical procedures, Xylazine provides analgesia and sedation however, it has substantial systemic effects on blood pressure and respiration. Ketamine does not effect the respiratory system but diminishes neurological responses due to NMDA-receptor antagonism (Hildebrandt et al., 2008). Both compounds are rapidly metabolized and fully excreted via the kidney and the liver. On the other hand, the specific pharmacological effects of α-chloralose are not fully elucidated but likely involve the GABAergic inhibitory system (Garrett and Gan, 1998). Due to its ability to preserve neurovascular coupling (neuronal activity and CBF responses) better than other anaesthetics, α-chloralose is traditionally used in Neuroscience (Lindauer et al., 1993; Bonvento et al., 1994; Austin et al., 2005; Sumiyoshi et al., 2012).

### Surgery

In our experimental set-up, the skull was glued to a metal plate using cyanoacrylate gel (Loctite Adhesives). One craniotomy was drilled for MCAO. Next, we drilled a second craniotomy with a diameter of _∼_4 mm, centered 0.5 mm behind and 3 mm to the right of the bregma, over the sensory barrel cortex region. After careful removal of the dura mater, anesthesia was switched to α- chloralose. The preparation was covered with 0.75% agarose in saline (type III-A, low EEO; Sigma-Aldrich), and then moistened with artificial cerebrospinal fluid (aCSF; NaCl 120 mM, KCl 2.8 mM, NaHCO_3_ 22 mM, CaCl_2_ 1.45 mM, Na_2_HPO_4_ 1 mM, MgCl_2_ 0.876 mM, glucose 2.55 mM; pH = 7.4) The craniotomy was partly covered with a glass coverslip that permitted electrode insertion for dye loading and neuronal activity recording (Fordsmann et al., 2018).

### Middle Cerebral Artery Occlusion

The mice were randomly assigned to either the sham control or MCAO groups. In the stroke groups, a small craniotomy was drilled over the distal part of the middle cerebral artery (MCA). The dura mater was removed, and the craniotomy was immediately moistened with aCSF and kept moistened during baseline laser speckle imaging (LSI) of cerebral blood flow (CBF). The MCA was electro-coagulated at the distal trunk using bipolar forceps coupled to an electrosurgical unit (Bipolar coagulator GN060; Aesculap). MCAO was confirmed by visual inspection through the surgical microscope and by LSI (Fordsmann et al., 2018). In the sham control group, a small craniotomy was drilled, but the MCA was left intact. Arterial blood gases were monitored and kept constant (PO_2_, 95–110 mmHg; PCO_2_, 35–40 mmHg; pH, 7.35–7.45; ABL 700Series Radiometer) during MCAO.

### Cerebral Blood Flow Monitoring by Laser Speckle Imaging

To validate our stroke model and identify ischemic areas in mice subjected to permanent MCAO, we monitored CBF with LSI (MoorFLPI; Moor Instruments) before, during, and at 10 min after MCAO, as well as in sham control mice, as previously described (Fordsmann et al., 2018). Images were acquired at 1 frame per second, and were processed offline using custom-made software (Python 2.7). In stroke mice, we defined the penumbra as an area where CBF was decreased to 20– 50% of pre-stroke values, and we defined an ischemic core by a drop to below 20% of pre-stroke values (Fordsmann et al., 2018). LSI recordings ensured the performance of two-photon microscopy imaging in areas that showed an initial CBF decrease corresponding to the ischemic penumbra. CBF was stable during LSI recordings in both adult sham and old sham controls, in which the MCA was untouched (Fordsmann et al., 2018). The ischemic core and penumbra sizes, and the relative decrease of CBF, were independent of age suggesting equivalent initial CBF reductions across all MCAO groups (Fordsmann et al., 2018).

### Whisker Pad Stimulation

Stimulations were performed 2–3 hours after stroke or sham intervention. To activate the mouse sensory barrel cortex, we percutaneously inserted a set of custom-made bipolar electrodes, and stimulated the contralateral ramus infraorbitalis of the trigeminal nerve (Norup Nielsen and Lauritzen, 2001). The cathode was positioned corresponding to the hiatus infraorbitalis, while the anode was inserted into the masticatory muscles. Stimulation was administered using an ISO-flex stimulation isolator (A.M.P.I., Israel), controlled by a sequencer file running within the Spike2 software version 7.02 (Cambridge Electronic Design, Cambridge, United Kingdom). Stimulation was applied for 0.1 ms with 1.5 mA in 15-s trains at 1, 2, and 5 Hz, with each train repeated twice.

### Electrophysiology

To record extracellular local field potentials (LFPs), we used a single-barreled glass micropipette filled with aCSF, along with the Ca^2+^-sensitive dye Oregon Green 488 BAPTA-1/AM (OGB) and the astrocyte-specific marker sulforhodamine 101 (SR101) (Lind et al., 2013). The Ag/AgCl ground electrode was positioned in the neck muscles while the recording electrode was inserted into the whisker barrel cortex (layer 2/3) and left untouched until the end of the experiments. All electrophysiological recordings were done in the ischemic penumbral area comprising layer 2/3 whisker barrel cortex. The signal was initially amplified using a differential amplifier (10× gain, 0.1–10,000 Hz bandwidth; DP-311 Warner Instruments). Then additional amplification was performed using the CyberAmp 380 (100× gain, 0.1–10,000 Hz bandwidth; Axon Instruments). The electrical signal was digitally sampled at a 5-kHz sampling rate using the 1401 mkII interface (Cambridge Electronic Design) connected to Spike2 software (Cambridge Electronic Design). For each stimulation train, we averaged the LFPs, and calculated the amplitudes of excitatory postsynaptic potentials as the difference between baseline and the first negative peak. Arterial blood gases were monitored and kept constant (PO_2_, 95–110 mmHg; PCO_2_, 35–40 mmHg; pH, 7.35–7.45; ABL 700Series Radiometer) during electrical recording.

### Two-Photon Microscopy

A glass micropipette (1–3 MΩ impedance; 2-μm tip; World Precision Instruments) was connected to a pneumatic injector (3–20 PSI, 10–90 s; Pneumatic Pump; World Precision Instruments). This set-up was used to load the somatosensory cortex with the Ca^2+^-sensitive dye Oregon Green Bapta (OGB-AM; Invitrogen) and the astrocyte-specific marker sulforhodamine (SR101; Sigma-Aldrich). OGB was solubilized in 20% pluronic acid and dimethylsulfoxide, and diluted to 0.8 mM in aCSF. SR101 was solubilized in methanol, and diluted to 0.005 mM in aCSF.

The same micropipette was also used for electrophysiological recordings. Care was taken to ensure calcium imaging within the ischemic penumbral area that comprised layer 2/3 whisker barrel cortex. LSI recordings ensured the performance of two-photon imaging in areas that showed an initial CBF decrease corresponding to the ischemic penumbra. We conducted *in vivo* imaging with two-photon microscopes using the Femto3D-RC (Femtonics, Hungary) and the SP5 multiphoton/confocal Laser Scanning Microscope (Leica, Germany) equipped with Ti:Sapphire lasers (Spectra-Physics, Sweden). Dye loading was performed under 5× magnification. Time-lapse movies were recorded using a 25× water-immersion objective (1.0 NA; Zeiss) and a 20× water-immersion objective (1.0 NA; Leica). Using an excitation wavelength of 900 nm, we recorded time-lapse movies with a frame size of 256 × 256 pixels (1.43 µm per pixel), and a sampling rate of 2–5 Hz (Fordsmann et al., 2018). Evoked Ca^2+^ activities were recorded at 120–180 min after the intervention. To minimize laser-induced artifacts and photo-toxicity, the laser power was kept between 20–40 mW.

### Image Analysis

Data were analyzed using a custom-built program written in Matlab. Regions of interest (ROIs) were selected using a modification of the pixel-of-interest-based analysis method (Lind et al., 2018). Rectangular ROIs were positioned around astrocytic somas, astrocytic processes, astrocytic end-feet, neuronal somas, and neuropil. ROIs were defined as either astrocytic or neuronal according to the SR101/OGB-AM double-staining pattern, cell morphology, and relation to blood vessels (Fordsmann et al., 2018). Large processes not visibly connected to a soma were considered astrocytic processes, while processes visibly connected to a soma were included as somatic ROIs. Astrocytic end-feet were identified based on the encircling of vessels. For each frame, we selected pixels showing intensities of 1.5 standard deviations (SD) above the mean intensity of the ROI. The intensities of these pixels were averaged, and then normalized to a 10-s baseline period just before stimulation onset, creating a time trace of ΔF/F_0_ for every ROI. These time traces were smoothed with a 2-sec moving average to avoid outlier values. An evoked Ca^2+^ response was defined as an intensity increase of ≥5% and of ≥2 SD from baseline, having a duration of ≥2.5 s, within 60 s after stimulation onset. Responsivity was defined as the fraction of ROIs responding to stimulation, and was reported as the % of responding ROIs among the total number of ROIs. For each ROI, response size was measured as the area under the curve (AUC) during the response, reported as AUC in ΔF/F_0_ × s.

### Experimental desing and statistical analysis

All statistical analyses were performed using RStudio version 1.0.136 (RStudio Team, 2016) with R version 3.3.2, (R Core Team, 2016) using ROIs as the experimental unit. Experimental groups were statistically compared to determine the effects of age, stroke, or pharmacological treatments. Differences in ROI responsivity were tested by multinomial logistic regression (MLR), with treatment and stimulation frequencies as explanatory variables, using the *nnet* package (Venables and Ripley, 2002). We tested the amplitude of excitatory postsynaptic potentials by performing linear mixed-effects modeling (LMER) with post-stroke time, treatment, and stimulation frequencies as fixed effects, and mice as random factors, using the *lme4* package (Bates et al., 2015). We also used the *lme4* program to assess the relationship between Ca^2+^ responsivities in different cellular compartments, with cellular compartment as a fixed effect and stimulation frequency as a random factor. Linear models (LM) were used to analyze the AUC of evoked Ca^2+^ increases, with treatment and stimulation frequencies as fixed effects. We used the Holm-Sidak post-hoc test to correct the *p* values obtained with MLR, LMER, and LM. To ensure normal distribution of residuals, the AUC for evoked Ca^2+^ increases was log-transformed before statistical analysis, and is presented as the back-transformed estimate ± the 95% confidence interval. The ROI % having evoked responses is presented as fraction ± standard error of probability. All other data are presented as mean ± standard error of the mean (SEM). The significance level was set to α = 0.05. Table 1 presents the numbers of mice, ROIs, and responses used for analysis.

**TABLE 1.** Numbers of mice, regions of Interest (ROIs), and responses for analyses of evoked Ca^2+^ responsivity and Ca^2+^ response Size. *ROI total* indicates the total number of regions of interest, with the number of mice used for analysis of evoked Ca^2+^ responsivity given in parentheses. *AUC (total)* indicates the total number of responses, with the number of mice used for analysis of evoked Ca^2+^ response size given in parentheses. ROI = regions of interest; AUC = area under the curve; MCAO = middle cerebral artery occlusion;

| Group | ROI (total) |  |  | AUC (total) |  |  |
| --- | --- | --- | --- | --- | --- | --- |
|  | 1 Hz | 2 Hz | 5 Hz | 1 Hz | 2 Hz | 5 Hz |
| Astrocyte soma |  |  |  |  |  |  |
| Adult sham | 100 (6) | 72 (5) | 79 (5) | 21 (6) | 22 (5) | 17 (5) |
| Adult MCAO | 59 (5) | 58 (5) | 61 (5) | 5 (3) | 7 (5) | 6 (2) |
| Old sham | 84 (5) | 149 (5) | 151 (5) | 24 (5) | 45 (5) | 41 (4) |
| Old MCAO | 62 (5) | 62 (5) | 57 (5) | 11 (5) | 13 (4) | 10 (3) |
| Astrocyte process |  |  |  |  |  |  |
| Adult sham | 92 (6) | 54 (5) | 67 (5) | 16 (5) | 16 (4) | 17 (4) |
| Adult MCAO | 76 (5) | 75 (5) | 77 (5) | 7 (3) | 4 (2) | 7 (5) |
| Old sham | 92 (5) | 162 (5) | 158 (5) | 28 (5) | 45 (5) | 31 (5) |
| Old MCAO | 104 (5) | 104 (5) | 95 (5) | 20 (4) | 15 (5) | 13 (4) |
| Astrocyte end-feet |  |  |  |  |  |  |
| Adult sham | 99 (6) | 66 (5) | 87 (5) | 15 (5) | 15 (5) | 12 (4) |
| Adult MCAO | 43 (5) | 42 (5) | 44 (5) | 2 (2) | 4 (3) | 1 (1) |
| Old sham | 100 (5) | 182 (5) | 182 (5) | 28 (5) | 65 (5) | 42 (4) |
| Old MCAO | 68 (5) | 67 (5) | 65 (5) | 8 (3) | 8 (5) | 6 (4) |
| Neuronal soma |  |  |  |  |  |  |
| Adult sham | 79 (6) | 67 (5) | 67 (5) | 23 (5) | 26 (3) | 15 (4) |
| Adult MCAO | 71 (5) | 71 (5) | 72 (5) | 7 (4) | 5 (4) | 5 (4) |
| Old sham | 49 (5) | 78 (5) | 79 (5) | 20 (5) | 24 (4) | 18 (3) |
| Old MCAO | 48 (5) | 48 (5) | 47 (5) | 6 (2) | 6 (3) | 7 (3) |
| Neuropil |  |  |  |  |  |  |
| Adult sham | 100 (6) | 80 (5) | 90 (5) | 28 (5) | 30 (4) | 25 (3) |
| Adult MCAO | 90 (5) | 90 (5) | 100 (5) | 2 (2) | - | 2 (2) |
| Old sham | 60 (5) | 100 (5) | 100 (5) | 49 (5) | 84 (5) | 62 (5) |
| Old MCAO | 100 (5) | 100 (5) | 90 (5) | - | - | 1 (1) |

## Results

### Focal Ischemia Abolishes Local Field Potentials

To determine how stroke affected synaptic excitation, we performed whisker pad stimulation and measured the LFP responses in the contralateral somatosensory cortex, which included the penumbra as indicated by LSI. MCAO abolished LFP responses in adult mice (*p* = 0.0065, LMER) and old mice (*p* = 0.0046, LMER; Figure 2 **and** Table 2).

**FIGURE 2:**
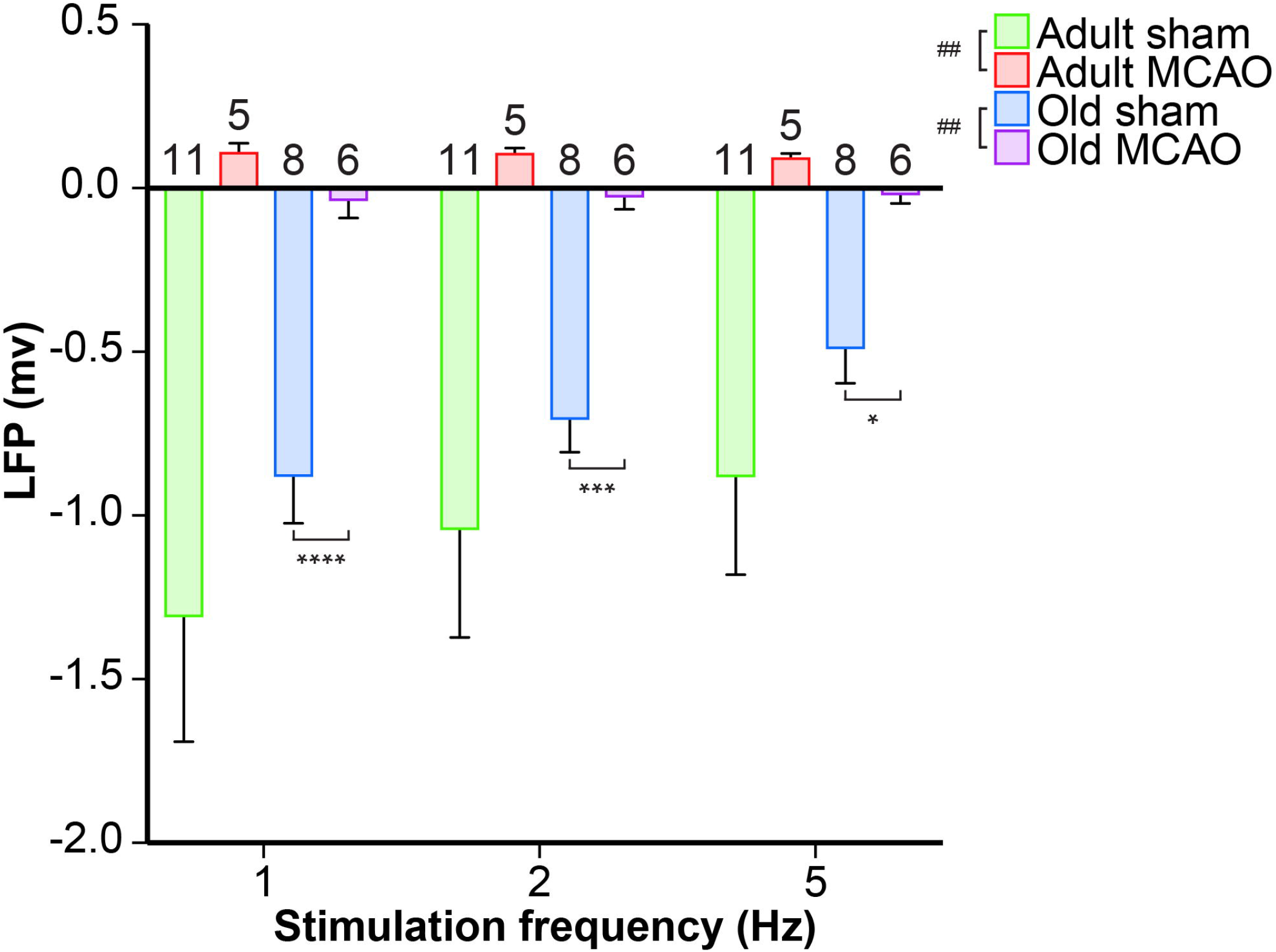
Stroke abolished local field potential (LFP) amplitudes in the penumbra of young and old mice. In old mice, middle cerebral artery occlusion (MCAO) diminished LFP amplitudes at all stimulation frequencies (1 Hz, *p* < 0.0001; 2 Hz, *p* = 0.0005; 5 Hz, *p* = 0.0207). Data are shown as mean ± standard error of mean. Data were analyzed using linear mixed-effects models with Holm-Sidak post-hoc tests. *Significant difference between treatments (i.e., experiment conditions). ^#^Significant difference in the interaction (treatment and stimulation frequencies) between experimental conditions. * and ^#^ in the legend indicate overall effects, while symbols over bars indicate group difference for an individual stimulation frequency. \**p* ≤ 0.05, \*\**p* ≤ 0.01, \*\*\**p* ≤ 0.001, \*\*\*\**p* ≤ 0.0001. The numbers of mice are presented over the bars.

**TABLE 2.** Amplitude of local field potentials. Data are presented as mean ± SEM in mV. MCAO = middle cerebral artery occlusion.

| Group | 1 Hz | 2 Hz | 5 Hz | Number of mice |
| --- | --- | --- | --- | --- |
| Adult sham | -1.31±0.38 | -1.04±0.33 | -0.88±0.30 | 11 |
| Adult MCAO | 0.11±0.03 | 0.11±0.02 | 0.09±0.01 | 5 |
| Old sham | -0.88±0.14 | -0.71±0.10 | -0.49±0.10 | 8 |
| Old MCAO | -0.04±0.05 | -0.03±0.04 | -0.02±0.03 | 6 |

### Stroke Strongly Reduces Evoked Neuronal Ca^2+^ Signals in the Ischemic Penumbra

Neurons and astrocytes generally respond to somatosensory stimulation with intensely increased Ca^2+^ in neuronal somas, neuropil, astrocytic somas, processes, and end-feet (Khennouf et al., 2016; Jessen et al., 2017; Lind et al., 2018; Stobart et al., 2018). Since stroke reduces spontaneous neuronal Ca^2+^ activity in adult but not aged mice, (Fordsmann et al., 2018) we next assessed how neurons in the electrically silent penumbra reacted to somatosensory stimulation during the early post-stroke period. Excitability was measured in terms of the ability of neurons and astrocytes to exhibit a Ca^2+^ response to synaptic input, presented as the response fraction, i.e., the number of responding ROIs relative to the overall number of ROIs. We also assessed response size, based on the AUC of the relative change in fluorescence ΔF/F_0_.

Healthy old and adult mice showed similar neuronal soma responsivity and Ca^2+^ response amplitudes. In adult mice under control conditions, somatosensory stimulation at 1, 2, and 5 Hz produced Ca^2+^ responses in 22–39% of neuronal somas. Relative to adult sham controls, stroke decreased this actively responding fraction by 66–82% (*p* < 0.0001, effect of treatment, MLR; Figure 3A–D). Additionally, stroke in adult mice reduced the Ca^2+^ response size by 34–88% (*p* = 0.0013, effect of treatment; LM, Figure 3E). In old mice, stroke reduced responsivity by 35–69% compared to age-matched sham controls (*p* = 0.0024, effect of treatment, MLR; Figure 3A–D), and decreased response size by 42–75% (*p* < 0.0002, effect of treatment, LM; Figure 3E).

**FIGURE 3:**
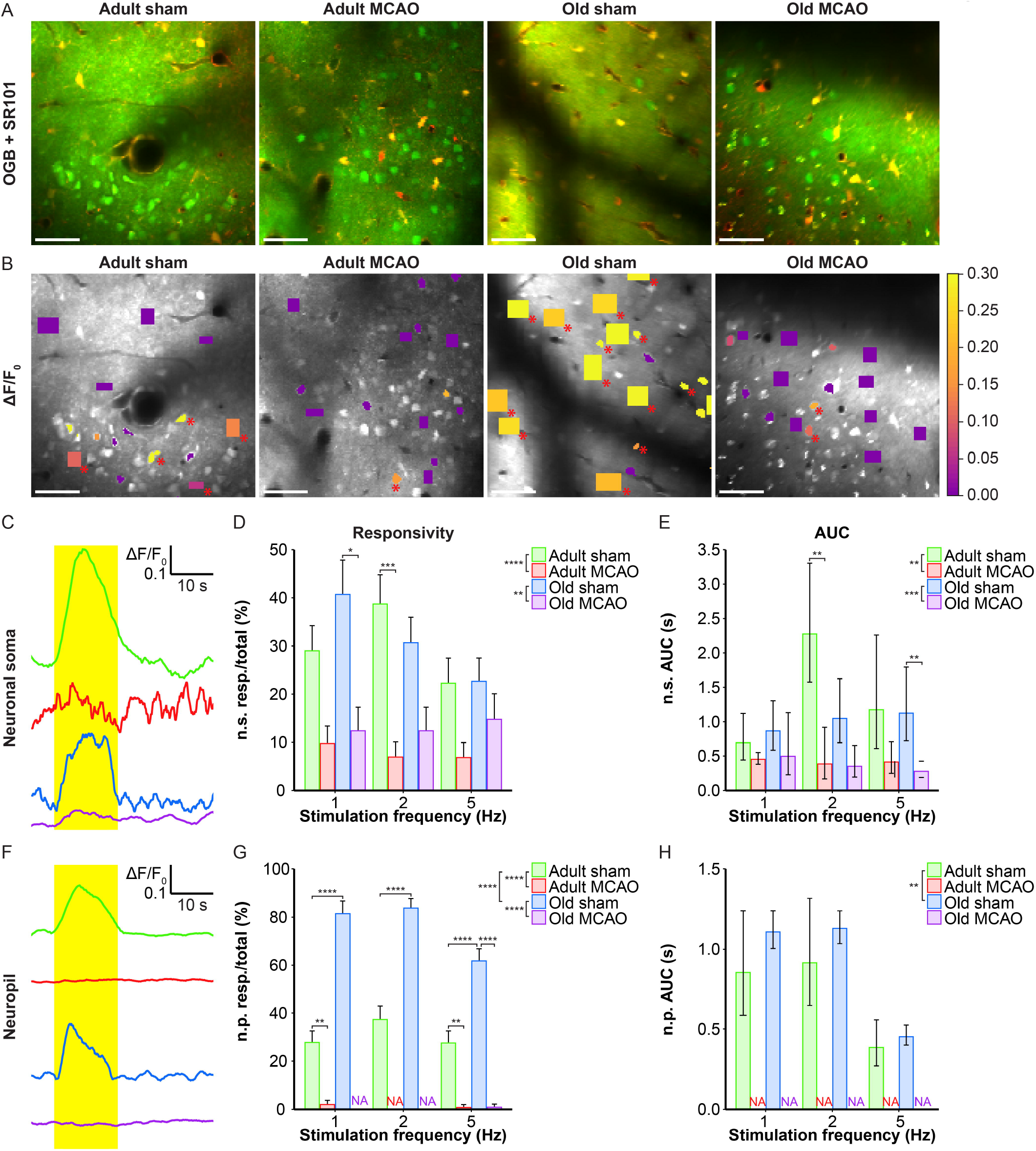
Stroke reduced the evoked Ca^2+^ signals in neuronal somas and neuropil in the penumbra. (**A, B**) Representative two-photon images recorded in adult and old mice after sham operation or middle cerebral artery occlusion (MCAO). (**A**) Green indicates Oregon Green 488 BAPTA-1/AM (OGB). Red indicates sulforhodamine 101 (SR101). (**B**) OGB is shown in gray. Mean ΔF/F_0_ during 2-Hz whisker stimulation is color-coded for neuronal somas (n.s.) and neuropil (n.p.). Red asterisk (*) indicates responsive regions of interest (ROIs). Scale bar = 50 μm. (**C, F**) Representative smoothed traces of ΔF/F_0_ from neuronal ROIs (**C**) and neuropil ROIs (**F**), upon stimulation at 2 Hz for 15 s, from adult sham (green), adult MCAO (red), old sham (blue), and old MCAO (purple) mice. (**D**) Compared to age-matched sham controls, stroke reduced the evoked neuronal soma responsivity in adult mice (2 Hz, *p* = 0.0006) and old mice (2 Hz, *p* = 0.0291). Responsivity is given as the fraction of responsive (resp.) ROIs relative to the total number of ROIs. (**E**) Compared to in age-matched sham controls, the evoked Ca^2+^ response area under the curve (AUC; ΔF/F_0_ × s) was reduced by 34–83% in the neuronal somas of adult and old mice after stroke. (**G**) Compared to age-matched sham controls, stroke reduced neuropil responsivity in adult mice (1 Hz, *p* = 0.0028; 5 Hz, *p* = 0.0080) and old mice (5 Hz, *p* < 0.0001). (**H**) Evoked neuropil Ca^2+^ response AUC (ΔF/F_0_ × s) during whisker stimulation. Neuropil responsivity and AUC increased with healthy aging. Values are given as the fraction ≤ standard error of probability for responsivity, and back-transformed estimates ± 95% confidence intervals for AUC. MCAO totally abolished evoked Ca^2+^ responses in neuropil; therefore, the AUC was not examined (NA). Responsivity was analyzed using multinomial logistic regression, and AUC using linear models—both with Holm-Sidak post-hoc tests. *Significant difference between treatments (i.e., experimental conditions). * in the legend indicates overall effects, while symbols over bars indicate the group difference for an individual stimulation frequency. \**p* ≤ 0.05, \*\**p* ≤ 0.01, \*\*\**p* ≤ 0.001, \*\*\*\**p* ≤ 0.0001.

We also examined the effect of stroke on Ca^2+^ responsivity and amplitude in the neuropil, i.e., the synaptically dense region between cell bodies that mainly comprises neuronal dendrites, unmyelinated axons, and fine astrocytic processes. Under control conditions, compared to adult mice, old mice showed enhanced neuropil responsivity (*p* < 0.0001, effect of treatment, MLR; Figure 3A, B, F, G), and larger neuropil Ca^2+^ response amplitudes (*p* = 0.0052, effect of treatment, LM; Figure 3H). MCAO abolished neuropil responsivity in adult mice (*p* < 0.0001, effect of treatment, MLR; Figure 3A, B, F, G) and in old mice (*p* < 0.0001, effect of treatment, MLR; Figure 3A, B, F, G). Overall, our data indicated that sensory stimulation evoked larger neuropil Ca^2+^ responses in old mice than adult mice under control conditions, and that stroke reduced neuronal soma responsivity and abolished neuropil Ca^2+^ responsivity in both adult and aged mice.

### Stroke Reduces Astrocytic Ca^2+^ Responses in Ischemic Penumbra in an Age-Dependent Manner

We next examined whether astrocytes in the ischemic penumbra responded to somatosensory stimulation after stroke. Under control conditions, adult and old mice showed similar evoked Ca^2+^ signaling in astrocytic somas, with a fraction of astrocytic somas exhibiting Ca^2+^ increases in response to sensory stimulation (Figure 4A, B **and** Figure 5B, C). After stroke, we observed critical age-dependent differences regarding evoked Ca^2+^ signals in astrocytic somas. In adult mice, stroke reduced the Ca^2+^ responsivity of astrocytic somas (*p* = 0.0029, effect of treatment, MLR; Figure 4A, B **and** Figure 5A, B), without altering the Ca^2+^ response amplitudes (Figure 5C). In contrast, among old mice, stroke did not reduce the responsivity or Ca^2+^ response amplitudes of astrocyte somas (*p* = 0.1098, MLR; and *p* = 0.1724, LM, respectively; Figure 5B, C). These data suggested that astrocytic soma excitability was more sensitive to stroke in adult mice than in aged mice.

**FIGURE 4:**
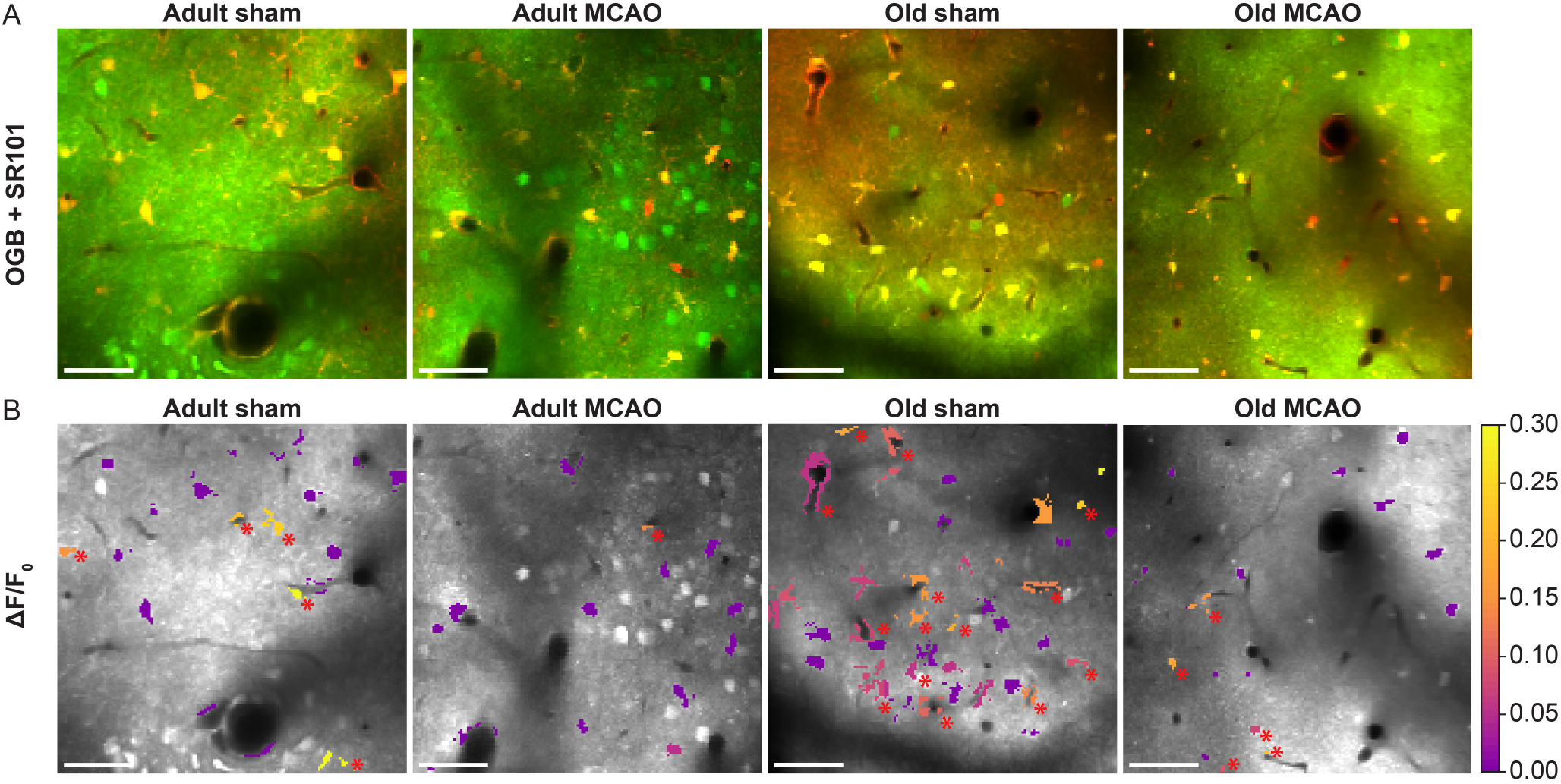
Stroke reduced evoked Ca^2+^ signals in astrocytic somas, processes, and end-feet. (**A, B**) Representative two-photon images recorded in adult and old mice after sham operation or middle cerebral artery occlusion (MCAO). (**A**) Green indicates Oregon Green 488 BAPTA-1/AM (OGB). Red indicates sulforhodamine 101 (SR101). (**B**) OGB is shown in gray. Mean ΔF/F_0_ during 2-Hz whisker stimulation for 15 s is color-coded for astrocytic somas (a.s.), astrocytic processes (a.p.), and astrocytic end-feet (a.e.-f.). Red asterisk (*) indicates responsive regions of interest. Scale bar = 50 μm.

**FIGURE 5:**
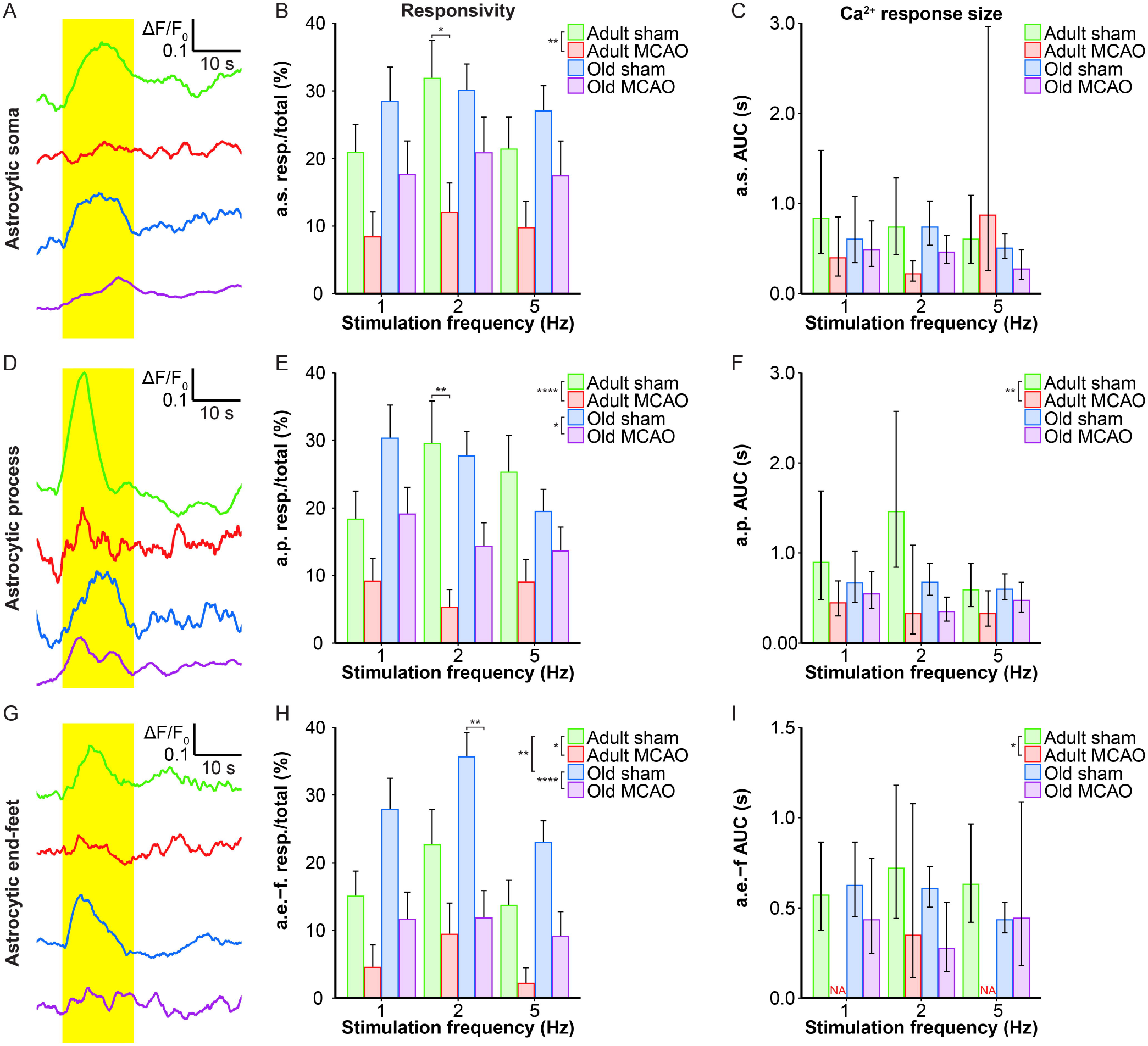
Quantification of the stroke-induced reduction of evoked astrocytic Ca^2+^ signals. (**A, D, G**) Representative smoothed traces of ΔF/F_0_ from astrocytic somas (a.s.) (**A**), astrocytic processes (a.p.) (**D**), and astrocytic end-feet (a.e.-f) (**G**), following stimulation at 2 Hz for 15 s, in adult sham (green), adult middle cerebral artery occlusion (MCAO; red), old sham (blue), and old MCAO (purple) mice. (**B, E, H**) Evoked Ca^2+^ responsivity of astrocytic somas, processes, and end-feet. Responsivity is shown as the fraction of responsive (resp.) regions of interest (ROIs) relative to the total number of ROIs. In adult mice, stroke decreased the responsivity of astrocytic somas, processes, and end-feet. In old mice, stroke only decreased the responsivity of astrocytic processes and end-feet (to a lesser degree than in adult mice). Stroke particularly decreased responsivity at the 2-Hz stimulation frequency in astrocytic somas (*p* = 0.0478; adult sham vs. adult MCAO), astrocytic processes (*p* = 0.0064; adult sham vs. adult MCAO), and astrocytic end-feet (*p* = 0.0048; old sham vs. old MCAO). (**C, F, I**) Size of the evoked Ca^2+^ response shown as AUC (ΔF/F_0_ × s) in astrocytic somas, astrocytic processes, and end-feet. In adult mice, stroke reduced evoked Ca^2+^ response size in astrocytic processes and end-feet. Values are given as fraction ≤ standard error of probability for responsivity, and as back-transformed estimates ± 95% confidence intervals for AUC. Stroke nearly abolished evoked Ca^2+^ responses in astrocytic end-feet at stimulation frequencies of 1 and 5 Hz; therefore, AUC was not examined (NA). Responsivity was analyzed using multinomial logistic regression, and AUC with linear models—and both with Holm-Sidak post-hoc tests. *Significant difference between treatments (i.e., experimental conditions). * placed in the legend indicates overall effects, while the symbol over bars indicates group differences for an individual stimulation frequency. \**p* ≤ 0.05, \*\**p* ≤ 0.01, \*\*\**p* ≤ 0.001, \*\*\*\**p* ≤ 0.0001.

Similar to astrocytic somas, under control conditions, sensory stimulation evoked similar Ca^2+^ signals in the astrocytic processes in the penumbra of both adult and old mice (Figure 5E, F). In adult mice, stroke reduced astrocytic process responsivity by 50–82% (*p* = 0.0001, effect of treatment, MLR; Figure 4A, B **and** Figure 5D, E) and reduced the Ca^2+^ response amplitude by 45– 77% (*p* = 0.0071, effect of treatment, LM; Figure 5F). Among old mice, stroke reduced the evoked Ca^2+^ responsivity in astrocytic processes by only 30–48% (*p* = 0.0128, effect of treatment, MLR; Figure 4A, B **and** Figure 5D, E) and did not alter the Ca^2+^ response amplitude (*p* = 0.1021, LM; Figure 5F). The results indicated that stimulation-induced Ca^2+^ signals in astrocytic processes were more susceptible to ischemia in adult mice than old mice.

Under control conditions, responsivity in the astrocytic end-feet was slightly higher in aged mice than in adult mice (*p* = 0.0015, effect of treatment, MLR; Figure 5H), while evoked Ca^2+^ response amplitude did not differ between groups (*p* = 0.9389, LM). Similar to our findings in astrocytic processes, stroke reduced responsivity in astrocytic end-feet by 58–84% in adult mice (*p* = 0.0324, effect of treatment, MLR; Figure 4A, B **and** Figure 5G**, H**) and by 58–67% in old mice (*p* < 0.0001, effect of treatment, MLR; Figure 4A, B **and** Figure 5G, H). Stroke reduced the evoked Ca^2+^ response size in astrocytic end-feet in adult mice (*p* = 0.0331; effect of treatment, LM; Figure 5I), but not in old mice (*p* = 0.1113, effect of treatment, LM; Figure 5I). Overall, stroke reduced the responsivity of Ca^2+^ responses in astrocytic end-feet in both adult and old mice, and reduced the Ca^2+^ response amplitudes in astrocytic end-feet only in adult mice.

### Relationship between Astrocytic and neuronal Ca^2+^ activity is disrupted by stroke

The decreased astrocytic responsivity after stroke may suggest a relationship between local evoked neuronal and astrocytic Ca^2+^ activity. To examine this relationship, we performed mixed models linear regression analysis with neuronal soma Ca^2+^ responsivity as the independent variable. Among adult sham controls, the Ca^2+^ responsivity of neuronal somas was significantly related to the responsivity of astrocytic processes, astrocytic end-feet, and neuropil (Table 3). In comparison, stroke in adult mice disrupted the relationship between Ca^2+^ responsivity of neuronal somas and Ca^2+^ responsivity of astrocytic processes and end-feet (Table 3). Among old sham controls, the Ca^2+^ responsivity of neuronal somas was significantly related to the responsivities of astrocytic somas and astrocytic processes. These relationships were also disrupted after stroke in old mice (Table 3). In conclusion, our data suggested that stroke disrupted the relationship between neuronal and astrocytic Ca^2+^ activity.

**TABLE 3.** Mixed models linear regression analysis of the relationships between Ca^2+^ responsivity in neuronal somas, neuropil, and astrocytes. Significant *p* values indicate a relationship between neuronal somas and the other cellular compartments. \**p* ≤ 0.05, \*\**p* ≤ 0.01, \*\*\**p* ≤ 0.001, \*\*\*\**p* ≤ 0.0001. *n* = number of recordings; MCAO = middle cerebral artery occlusion.

| Group | Astrocytic somas |  | Astrocytic processes |  | Astrocytic end-feet |  | Neuropil |  |
| --- | --- | --- | --- | --- | --- | --- | --- | --- |
|  | *p* | *n* | *p* | *n* | *p* | *n* | *p* | *n* |
| Adult sham | 0.2000 | 24 | 0.0038** | 24 | 0.0001**** | 24 | <0.0001**** | 24 |
| Adult MCAO | 0.2011 | 29 | 0.4133 | 29 | 0.2892 | 29 | 0.6211 | 27 |
| Old sham | 0.0113* | 23 | <0.0001**** | 23 | 0.1546 | 23 | 0.0063** | 23 |
| Old MCAO | 0.8884 | 29 | 0.2226 | 29 | 0.0251* | 29 | 0.5518 | 29 |

## Discussion

Functional studies in rodents provide evidence that administration of mild sensory stimulation within a critical time window of a few hours after stroke can protect the cortex from impeding ischemic injury, but the mechanisms are incompletely understood, (Lay et al., 2011; 2012; Frostig et al., 2013; Hancock et al., 2013; Lay and Frostig, 2014; von Bornstadt et al., 2018) and additional studies are needed to clarify these findings (Baron, 2018). Here we report that in early stroke, the astrocytic Ca^2+^ signals in the penumbra were moderately preserved, while electrical potentials and neuronal Ca^2+^ responses were almost silenced. We speculate that astrocyte activity in the electrically silent penumbra may mechanistically contribute to functional recovery after ischemic stroke.

In a recent publication, we demonstrated increased *spontaneous* astrocytic Ca^2+^ activity and stroke-resistant *spontaneous* neuronal Ca^2+^ activity in the penumbra of old mice, while neuronal and astrocytic spontaneous Ca^2+^ activity were reduced in adult mice (Fordsmann et al., 2018). In our present study, we found that electrical silence did not exclude stimulation-evoked Ca^2+^ signaling in the penumbra. These results suggest a mismatch between Ca^2+^ signaling and electrophysiological signals in the penumbra. Notably, astrocytic Ca^2+^ signals were better preserved in the penumbra of old mice than adult mice. The exact mechanism underlying these age-dependent effects is unknown, and it remains to be determined whether penumbral astrocytic Ca^2+^ signals in aged mice are adaptive, beneficial, or damaging. Stroke predominantly occurs in elderly people, and age strongly influences patient outcomes after stroke; however, most preclinical studies have been performed in young animals, and thus have less translational potential than data from an aged experimental group (Liu et al., 2009; Chen et al., 2010; Manwani and McCullough, 2011; Selvamani et al., 2014). Astrocytes may positively or negatively influence tissue survival depending on their phenotype, which dramatically changes during a life course (Bhat et al., 2012). In old mice, astrocytes transform to a reactive neuroinflammatory phenotype that produces complement components, and releases toxic factors that kill neurons and oligodendrocytes in the uninjured brain (Clarke et al., 2018). During ischemia, this astrocyte phenotype is likely to change to also express ischemia-induced genes (Clarke et al., 2018). It may be possible to enhance neuroprotection by blocking astrocyte conversion to the inflammatory or ischemic phenotype, (Yun et al., 2018) but additional studies are needed to examine this possibility.

Few prior studies have examined astrocytic Ca^2+^ signaling *in vivo* in the penumbra. Reports describe increased astrocytic Ca^2+^ in association with large *spontaneous* Ca^2+^ waves, reminiscent of cortical spreading depolarization waves, during the first 1–1.5 hours after vascular occlusion (Ding et al., 2009; Rakers and Petzold, 2017). Exposure to mild sensory stimulation may also elicit cortical depolarization waves at later times in susceptible cortical regions, (von Bornstadt et al., 2015) but this was not observed in our present study. Even in the absence of cortical spreading depolarization waves, the ischemic penumbra is electrically silenced due to low blood flow, (Astrup et al., 1977) as was evident in our adult and old mice. Consistent with previous findings, our present results support that neurons are more sensitive to ischemia than astrocytes, since stroke abolished all types of neuronal Ca^2+^ activity in both young and old mice, but did not abolish astrocyte activity (Duffy and MacVicar, 1996; Brown, 2004).

Studies using genetically encoded Ca^2+^ indicators have detected spontaneous and evoked Ca^2+^ signals in fine processes around synapses, which exhibit spatial and temporal variation (Shigetomi et al., 2016; Verkhratsky and Nedergaard, 2018). The exact function of Ca^2+^ signals in each of these microdomains is still being investigated. In our present study, we did not assess Ca^2+^ signals from finer astrocytic branches because these branches cannot be labeled with organic dyes. On the other hand, compared to genetically encoded dyes, organic dyes label a greater portion of astrocytes, and our Ca^2+^ signals mainly stemmed from astrocyte somas, thick processes, and end-feet. The low SR101 concentration (≤50 μM) used in our study did not induce seizure activity, which has been observed in other studies (Kang et al., 2010; Rasmussen et al., 2016). In our sham controls, the responsivity of astrocyte somas, processes, and end-feet to stimulation was similar across structures, consistent with previous findings (Gu et al., 2018).

We identified a significant correlation between evoked neuronal and astrocytic Ca^2+^ signals in control mice, which was disrupted after stroke in both adult and old mice. These findings suggest that stroke interrupts neuronal control over astrocytic Ca^2+^ signaling, Ischemia-induced energy depletion may result in the release of glutamate, GABA, and ATP, which may affect Ca^2+^ signaling and interactions between astrocytes and neurons in the penumbra (Fellin et al., 2004; Wang et al., 2006; Attwell et al., 2010).

In summary, here we used *in vivo* two-photon microscopy to measure the cytosolic Ca^2+^ responses evoked by physiological stimulation in the penumbra of adult and old mice. We report that moderately preserved astrocytic Ca^2+^ responses may modulate the neuroprotective effect of sensory stimulation in early stroke. Moreover, astrocytic Ca^2+^ signals were better preserved in old than in young adult mice. Modulation of astrocyte activity may be considered in future neuroprotective strategies.

## Acknowledgements

We thank Zindy Raida, Anders Hay-Schmidt, and Micael Lønstrup for surgical assistance; and Barbara L. Lind, Krzysztof Kucharz, and Claus Mathiesen for helpful discussions. This study was supported by the University of Copenhagen (R40-A3311), the Lundbeck Foundation (R126- A12462), the NORDEA Foundation Grant to the Center for Healthy Aging (315600), the NOVO-Nordisk Foundation (NNF15OC0017366), the Russian Science Foundation (17-74-20089), and the Danish Council for Independent Research | Medical Sciences (0602-01965B FSS).

## Author Contributions Statement

RM, JF, and ML contributed to the study conception and design; RM, JF, CC, and KT contributed to data acquisition and analysis; RM, JF, KT, and ML contributed to drafting the text and preparing the figures.

